# *Burkholderia cenocepacia* physiology and molecular adaptations to the acidic pH of the CF nutritional environment

**DOI:** 10.64898/2026.08.03.742460

**Authors:** L. D. Morales, B. K. Dhillon, J. C. Grigg, A. Saraph, L. D. Eltis, R. E. W. Hancock, M. E. P. Murphy

## Abstract

*Burkholderia cenocepacia* is an opportunistic pathogen associated with increased disease severity and mortality in cystic fibrosis (CF) patients. We have previously shown that elevated iron and acidic pH in the CF nutritional environment increases *B. cenocepacia* growth rate and decreases its susceptibility to some of the antimicrobials used clinically to treat CF infections. Here, we aimed to characterize *B. cenocepacia* physiology and its molecular response under acidic pH and increased zinc and iron concentrations using a modified synthetic CF sputum media (SCFM-FeZn). By investigating *B. cenocepacia* internal pH homeostasis, we found that it maintains a neutral internal pH when exposed to mildly acidic media at pH 5.5. We also assessed the effect of *B. cenocepacia* growth on the pH of SCFM-FeZn. When cultured at pH 6.8, *B. cenocepacia* maintained a media pH of ∼6.5. In contrast, when the culture pH value was initially 5.5, it increased to 6.5 during growth. Using comparative transcriptomics and metabolomics analysis, we identified 990 differentially expressed genes, and 23 differentially abundant metabolites in supernatants at acidic compared to neutral pH. Some of these genes and metabolites were involved in aromatic amino acid metabolism including the upregulated *trpE* gene, encoding a tryptophan biosynthetic enzyme. A tryptophan auxotrophic *trpE* deletion strain grew slower in SCFM-FeZn. Overall, this work identifies mechanisms involved in *B. cenocepacia* adaptation to acidic pH under conditions to model the CF nutritional environment. Some of these mechanisms are also associated with pathogenicity and virulence.

**Importance:** Pathogenic bacteria can be exposed to acidic pH inside and outside the host. Their ability to adapt to pH fluctuations contributes to success in host colonization. B. cenocepacia can grow at acidic pH (∼3.5) and has been recovered from intracellular acidic compartments of amoebas and macrophages. Adaptation to acidic pH depends on molecular mechanisms that maintain a near optimal pH inside the cell for the function of vital processes. A few mechanisms that contribute to its adaptation to acidic pH have been described, but not in conditions reflecting the CF nutritional environment. Here, we identified multiple differentially-regulated systems that are associated with bacterial susceptibility to antimicrobials and pathogenesis. This research provides a better understanding of the role of acidic pH on B. cenocepacia physiology in the CF nutritional context and highlights possible systems that should be further characterized.

## Introduction

Cystic Fibrosis (CF) is a genetic disease caused by mutations in the human cystic fibrosis transmembrane conductance regulator (CFTR) gene. These mutations lead to accumulation of viscous and sticky mucus in the ducts and airways of CF patients and facilitate the development of chronic bacterial infections (1,2). Malfunction of CFTR impacts the pH homeostasis of the lung. CF airways are more acidic when compared to normal individuals and direct measurements of CF sputum samples show pH ranging from 2.9 to 6.5 compared to the neutral pH of healthy sputum (3–5). Infection is itself a major contributor to acidification. For example, CF pathogens like *Pseudomonas aeruginosa* and *Staphylococcus aureus* produce acids that contribute to the localized reduction in pH (6–8). Also, in CF, acidic pH is associated with lung infection severity since patients with infective exacerbations have lower airway pH than do those that are clinically stable (9). During bacterial infections, hypoxia and anaerobic glycolysis cause the accumulation of metabolites such as lactic acid, which when combined with host metabolites contribute to local acidosis (10,11). Acidogenesis and aciduricity are bacterial mechanisms to outcompete acid-sensitive bacteria and decrease microbial diversity (12).

Bacterial success in colonization is associated in part with ability to cope with pH fluctuations inside the host. Bacteria have multiple molecular mechanisms to adapt to environmental pH gradients. Response to acidic stress has been extensively studied in gastrointestinal pathogens such as *E. coli*, *S. enterica* and *H. pylori* (13–15). Bacterial adaptation to acidic pH are classified based on the following mechanisms: enhancement of proton pumping, alteration of cell membranes, protection and repair mechanisms, and sequestration of intracellular protons (16,17). Some of the mechanisms triggered by external pH acidification are associated with pathogenesis, host colonization, and antimicrobial resistance in these species. For example, membrane modification triggered by acidic environmental pH in *S. enterica* and *Vibrio fischeri* increases resistance to cationic antimicrobials (18,19). Similarly, comparative transcriptomics of *Brucella melitensis* grown at acidic and neutral pH show the upregulation of genes associated with intracellular survival, virulence and persistence (20).

*Burkholderia cenocepacia* is a highly transmissible opportunistic pathogen and causes severe lung infections associated with increased morbidity and mortality in CF patients (21). *Burkholderia* species are abundant in mildly acidic soils and *B. cenocepacia,* in particular, grows at pH values as low as 3.5 (22). Interestingly, the molecular mechanisms for coping with acidic pH also contribute to persistence in the host. Specifically, *B. cenocepacia* cells survive inside acid compartments of amoebas, persist in macrophages, and delay phagosomal acidification (23,24). Hopanoid production is a mechanism associated with growth under acidic pH in *B. cenocepacia* (25). However, molecular mechanisms to cope with acidic pH in the CF nutritional environment are unknown.

Previously, we modified synthetic CF media (SCFM) by decreasing the pH and increasing the iron and zinc content to better mimic the physiological conditions encountered by *B. cenocepacia* in the lung (26). *B. cenocepacia* grew at higher rates in this media, termed SCFM- FeZn, and many antibiotics currently used to treat CF infections failed to inhibit growth in this media. Here, we characterized *B. cenocepacia* physiology growing in SCFM-FeZn and used this media to identify molecular mechanisms potentially expressed to adapt to the acidic pH in the CF nutritional environment. Our aim is to provide a better understanding of *B. cenocepacia* physiology and molecular mechanisms triggered by acidic pH that could be associated with pathogenesis, virulence and antimicrobial resistance.

## Results

### B. cenocepacia increased the pH of its extracellular environment during growth

First, we assessed the impact of *B. cenocepacia* on its extracellular environment by assessing the pH of SCFM-FeZn during bacterial growth. *B. cenocepacia* cultures were grown for 24 h in SCFM-FeZn initially adjusted to pH 5.50 or 6.80 and supplemented with a fluorescent pH indicator, BCECF (Figure 1). To control for the effect of medium buffering, SCFM-FeZn with low buffering capacity (10 mM MOPS and 10 mM MES) and high buffering capacity (25 mM MOPS and 25 mM MES) were used. A calibration curve in SCFM-FeZn was used to transform BCECF fluorescence readings into pH values (Supplemental Figure S1). At a starting pH of 6.50, the media pH remained close to neutrality in media with either buffering capacity (Figure 1A). We observed a slight decrease in the media pH during the first eight hours of bacterial growth, followed by an increase to 6.66 ± 0.06, and 6.49 ± 0.09, for media with high and low buffering capacity, respectively. At a starting pH of 5.30, the media pH increased over time and reached values of 5.77 ± 0.01 and 6.47 ± 0.04, for media with high and low buffering capacity, respectively (Figure 1B). As a control, the fluorescence of uninoculated media was measured over time. Interestingly, when culturing *B. cenocepacia* at acidic pH with low buffering capacity, we observed an increase in neighboring wells of the medium control; this is consistent with the possibility that *B. cenocepacia* metabolites that affected the pH of the media were volatile (data not shown). *B. cenocepacia* cells cultured at acidic pH grew at a faster rate when compared to those at neutral pH as previously reported (26), and the media buffering capacity did not impact on bacterial growth. Our data indicates that at acidic pH, *B. cenocepacia* metabolism increases the pH of a nutritional environment equivalent to that in CF.

**Figure 1.**
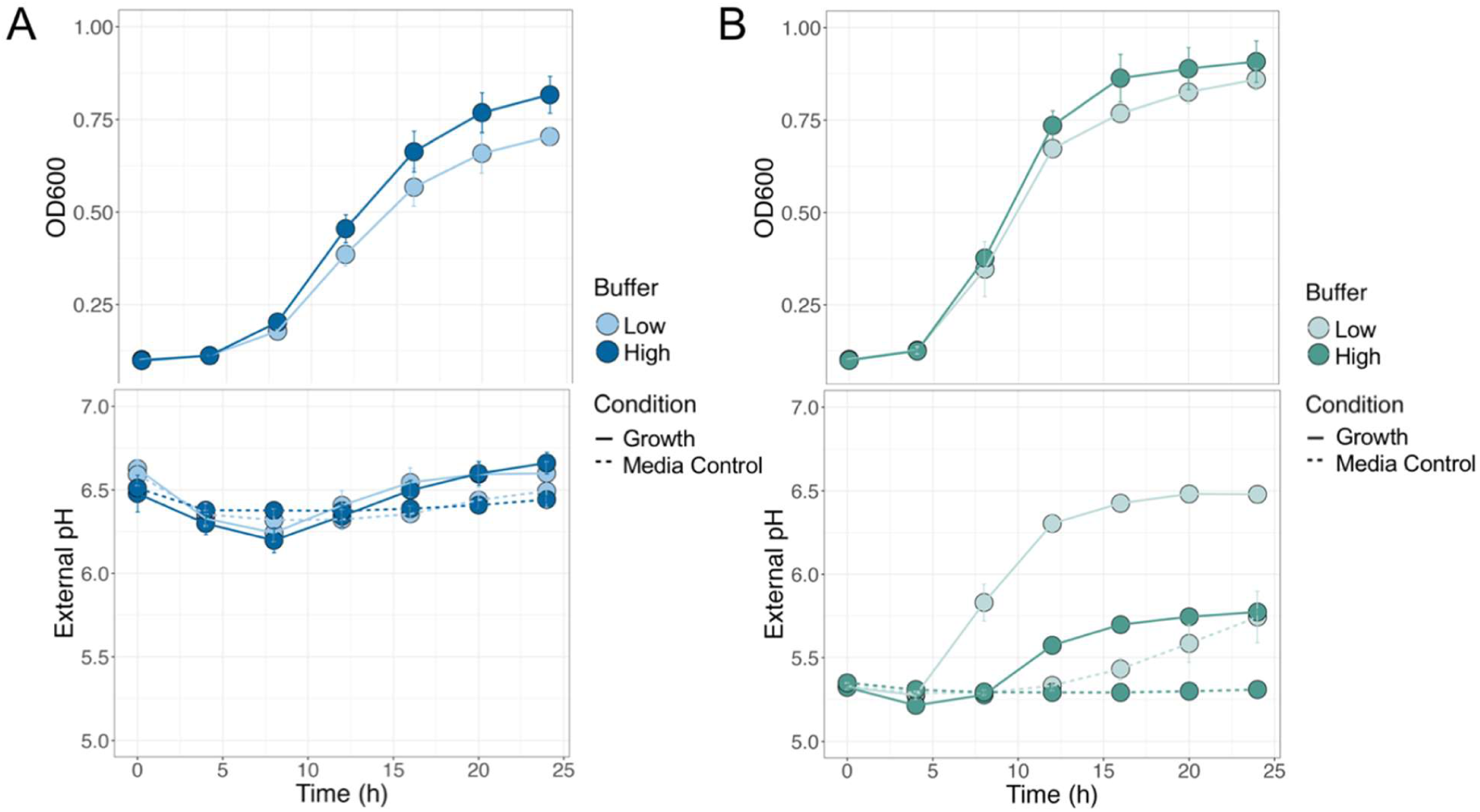
Impact of *B. cenocepacia* growth on the pH of the SCFM-FeZn. *B. cenocepacia* K56-2 was inoculated in SCFM-FeZn media at pH 6.80 **(A)** and 5.50 **(B)** supplemented with 0.1 μM BCECF. Bacterial growth and media pH was monitored for 24 hours by measuring optical density (OD_600_) and fluorescence every four hours. A BCECF calibration curve was used to transform the fluorescence intensities into pH values. SCFM-FeZn with two buffering capacities was used: Low (10 mM MOPS and 10 mM MES) and high (25 mM MOPS and 25 mM MES). Dotted line represents the media control with no *B. cenocepacia* K56-2 cells inoculated. Measurements were taken with three replicates on two independent days, n= 6. Error bars represent the standard deviation between individual measurements (SD).

### B. cenocepacia maintained a neutral intracellular pH after extracellular pH acidification

The stability and function of the biological molecules within the bacterial cell is dependent on the intracellular pH (37). Bacteria employ different strategies to maintain a narrow intracellular pH (38). We characterized the intracellular pH of *B. cenocepacia* by using a pH sensitive derivative of a green fluorescent protein, PHP, that was optimized for expression in *Pseudomonas* sp. (27,39). The PHP DNA sequence was inserted into a pMLBAD vector and transformed into *B. cenocepacia* K56-2. To identify pH-dependent fluorescence in *B. cenocepacia,* PHP-expressing cells were suspended in M9 media at pH values ranging from 5.0 to 8.5. Proton gradient uncouplers (50 mM sodium benzoate and 50 mM methylamine HCl) were added to the cell suspensions to collapse the transmembrane proton gradient and allow the intracellular pH to equilibrate with the extracellular pH. The pH-dependent excitation maxima of PHP were found to be 400 and 478 nm (Supplemental Figure S2A) in *B. cenocepacia*. A calibration curve was constructed from the ratio of the fluorescence at 400 / 478 nm (Supplemental Figure S2B).

To assess how *B. cenocepacia* intracellular pH changes with altered extracellular pH, cells were resuspended in M9 media at pH 5.50, 7.00, and 8.50. Proton uncouplers were added to some cell suspensions to collapse the ΔpH. *B. cenocepacia* intracellular pH measurements in M9 are shown in Figure 2A. The observed intracellular pH in M9 was 7.51 ± 0.19 at neutral pH was, 7.11 ± 0.08 at acidic pH and 8.10 ± 0.09 M9 at alkaline pH. The intracellular pH observed under the three conditions remained approximately constant for 10 min of measurement (data not shown).

**Figure 2.**
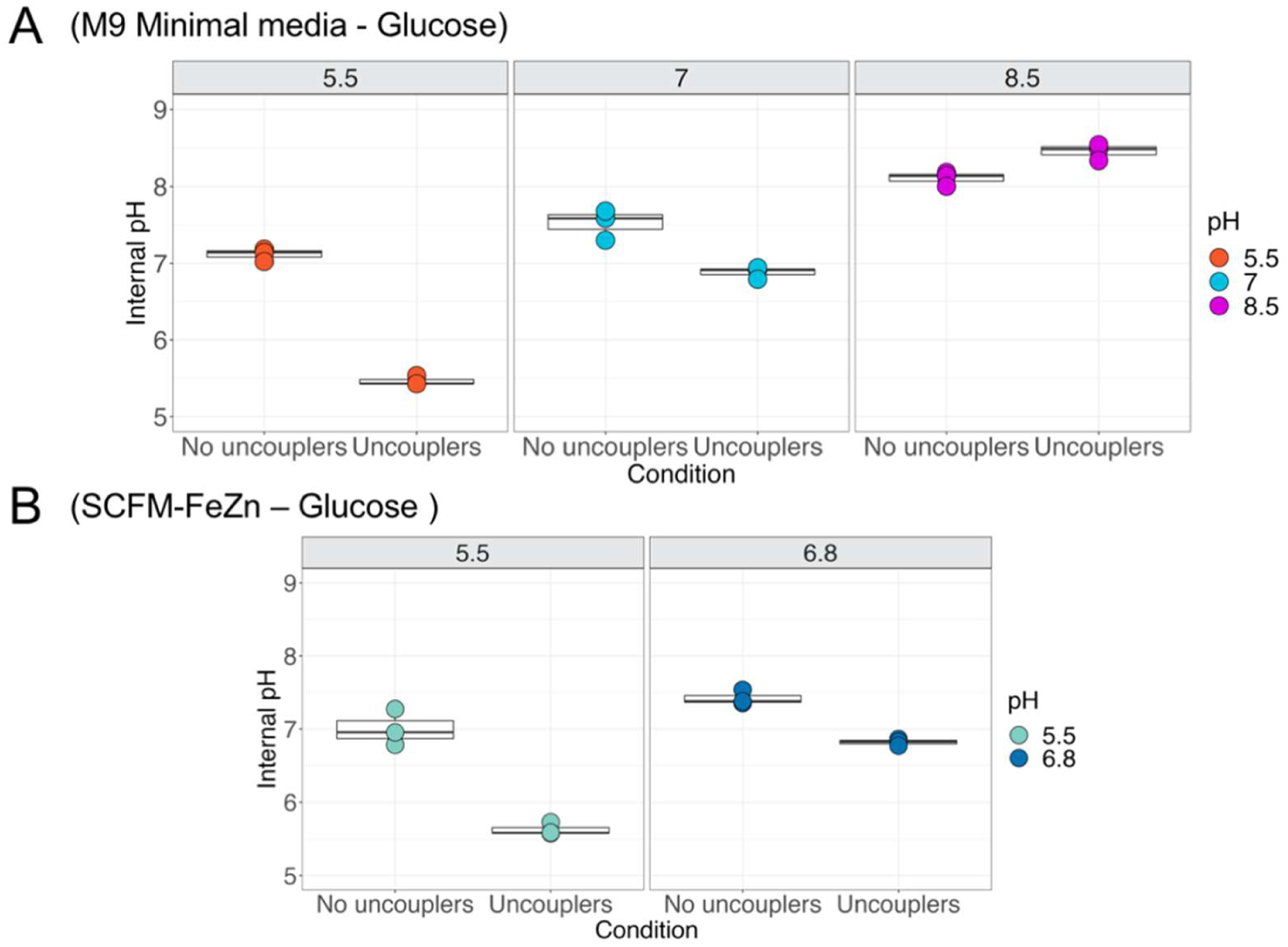
*B. cenocepacia* maintains a neutral internal pH after external pH acidification. Internal pH was monitored for cells in M9 mimimal media **(A)** and SCFM-FeZn **(B)**. Internal pH was determined by measuring fluorescence of *B. cenocepacia* K56-2 expressing PHP after two minutes of incubation at each pH value (λ_Em_: 515 nm, λ_Ex_=400 and 478 nm). As media controls, some of the cells were resuspended in media supplemented uncouplers. Internal pH was measured on three biological replicates (n= 3).

Molecules such as weak organic acids can diffuse across the plasma membrane and neutralize protons intracellularly reducing the intracellular pH (40). To characterize the role of the nutritional environment on *B. cenocepacia* intracellular pH homeostasis, PHP-expressing cells were suspended in SCFM-FeZn at pH 5.50 and 6.80 (Figure 2B). The intracellular pH of cells in SCFM-FeZn behaved similar to that of M9. The observed intracellular pH was 7.42 ± 0.09 for cells resuspended in the medium at pH 6.80, and 7.00 ± 0.24 in medium at pH 5.50. Our data suggest that *B. cenocepacia* has mechanisms to maintain a near neutral internal pH when exposed to acidic conditions but not to alkaline conditions.

### B. cenocepacia comparative transcriptomics and metabolomics in the SCFM-FeZn

The mechanisms involved in *Burkholderia* sp. acid tolerance have not been described. To elucidate the mechanism for adaptation to acidic pH in a model of the CF nutritional environment, we conducted transcriptomics and culture supernatant metabolomics of *B. cenocepacia* grown in the SCFM-FeZn at acidic and neutral pH (Supplemental Figure S3). Oxygen profiling of CF sputum samples revealed that oxygen is depleted a few millimeters below the sample’s surface (3). Thus, for transcriptomics and supernatant metabolomics analysis, cells were grown at low aeration (50 rpm) to approximate the decreased oxygen levels.

The transcriptomes of *B. cenocepacia* grown in SCFM-FeZn media at pH 5.50 and 6.80 were compared to identify differentially expressed (DE) genes at acidic pH. In a principal component analysis (PCA) of the transcriptomes, samples are separated along PC1 (81% of the variance) according to the media pH (Figure 3A). A total of 990 differentially expressed genes (≥ 2-fold change and adjusted *p*-value < 0.05) (Figure 3B) were identified, of which 597 were upregulated and 393 were downregulated at acidic pH (see Supplemental Table S1). Examining the list of genes, it was evident that growth at low pH is a highly regulated process with 83 one-component regulators being upregulated by 2- to 246-fold and 10 being downregulated by 2- to 3.5-fold. In addition, 9 two-component response regulators and 2 sensor kinases were upregulated and 4 response regulators and 1 sensor kinase were down regulated. One sigma factor also showed a 2.3-fold increase in expression. This indicates a likely regulon responding to mildly acidic conditions. Hints as to the function of at least one set of genes was found by the presence of a response regulator/sensor kinase pair of genes upstream of a phosphate utilization operon comprising phoR-phoB-phoU-pstBAS. This might relate to the state of phosphate at different pH values with a pKa of 7.2 for the transition from monobasic to dibasic phosphate (Supplemental Table S1).

**Figure 3.**
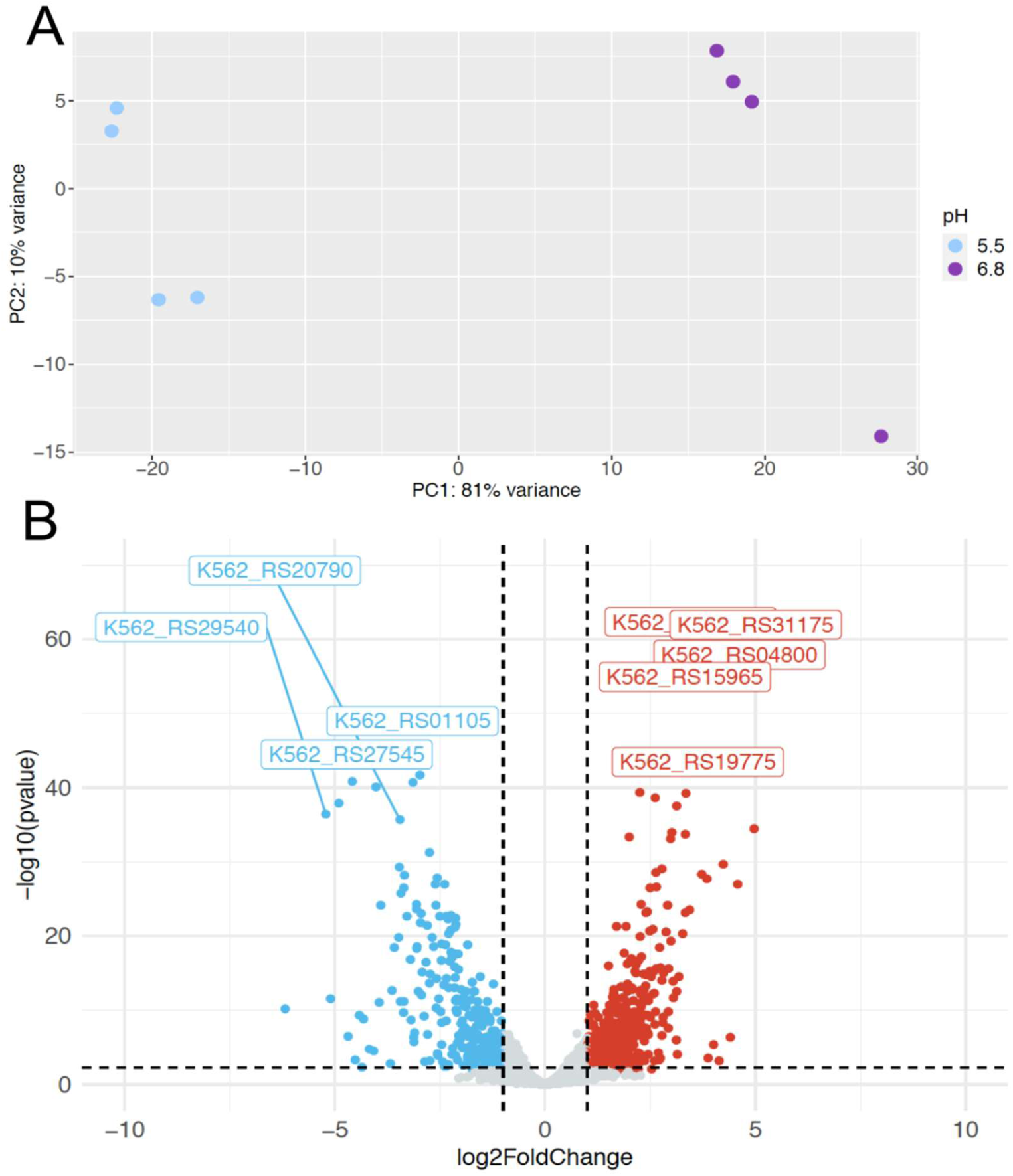
Comparison of the transcriptomes of *B. cenocepacia* K56-2 grown in the SCFM-FeZn at acidic and neutral pH. **(A)** A PCA plot to visually inspect the variation in gene expression between samples. Samples from cells grown at pH 5.5 and pH 6.8 are shown in blue and purple, respectively **(B)** A volcano plot to idenitify differentially expressed genes. Cutoff for selected DE genes was expression fold change >2 and *p*-value <0.05. Downregulated genes are shown in blue and uregulated genes are shown in red.

There is a modest literature around the effects of acidic stress on bacterial gene expression (41). Key effects noted are those for (a) homeostasis and defense including upregulation of proton pumps (such as ATPases) and amino acid degradation enzymes (e.g. glutamate decarboxylase) to neutralize intracellular pH shifts (41). and (b) virulence factors and flagella motility (42). Amongst the dysregulated genes we observed for category (a) a 3.6-fold upregulated AAA family ATPase (RS02790), and 3 downregulated ATPases, including *kdpABC*, as well as amino acid degrading enzymes such as RS30175 encoding 4- aminobutyrate-2-oxoglutarate transaminase. Virulence and motility factors in category (b) included 3 modestly upregulated toxin-antitoxin systems and flagella regulators *flhCD*, while a Type III secretion system Sct (which delivers toxins directly to host cells) had multiple components downregulated (Supplemental Table S1).

Pathway enrichment analysis was performed on the DE genes list (Supplemental Table S1) using KEGGREST which assigns genes to biochemical pathways for genomes summarized in the Kyoto Encyclopedia of Genes (KEGG). Since *B. cenocepacia* K56-2 was not available in KEGG, differentially expressed genes were assigned to pathways in the genome of strain *B. cenocepacia* J2315 for genes whose protein accession numbers were identical. KEGGREST analysis revealed ten pathways that were significantly enriched (*p*-value < 0.05) (Table 1). Of these ten, two were involved in amino acid metabolism, and three in carbohydrate metabolism. A majority of the genes assigned to phenylalanine metabolism are part of the phenylacetic acid pathway, and most of the identified genes annotated as in carbohydrate metabolism are involved in cepacian biosynthesis.

**Table 1.** *B. cenocepacia* K56-2 pathway enrichment analysis using KEGGREST. D. E genes with identical proteins IDs to *B. cenocepacia* J2315 were used to apply a Wilcoxon rank-sum test to test for enrichment of each pathway.

| Pathway name | Hits | Pathway Size | Genes |
| --- | --- | --- | --- |
| Phenylalanine metabolism | 12 | 53 | K562_RS01095, K562_RS01105, K562_RS01110, K562_RS15375, K562_RS25815, K562_RS25820, K562_RS02100, K562_RS02095, K562_RS02105, K562_RS01100, K562_RS01115, K562_RS14005 |
| Pentose phosphate pathway | 3 | 34 | K562_RS14635, K562_RS09660, K562_RS30300 |
| O-Antigen nucleotide sugar biosynthesis | 5 | 20 | K562_RS21500, K562_RS21490, K562_RS22290, K562_RS22255, K562_RS22250 |
| Amino sugar and nucleotide sugar metabolism | 8 | 47 | K562_RS21500, K562_RS21490, K562_RS22290, K562_RS22255, K562_RS22250, K562_RS27440, K562_RS03175, K562_RS05330 |
| Galactose metabolism | 2 | 20 | K562_RS04775, K562_RS22290 |
| Glutathione metabolism | 3 | 47 | K562_RS19775, K562_RS14635, K562_RS02840 |
| Bacterial chemotaxis | 5 | 40 | K562_RS23195, K562_RS27375, K562_RS21125, K562_RS23215, K562_RS24730 |
| Fructose and mannose metabolism | 7 | 28 | K562_RS21490, K562_RS22255, K562_RS05180, K562_RS05115, K562_RS22250, K562_RS31225, K562_RS03175 |
| Lysine biosynthesis | 4 | 20 | K562_RS31220, K562_RS03315, K562_RS22165, K562_RS19825 |
| Monobactam biosynthesis | 4 | 12 | K562_RS31220, K562_RS03315, K562_RS22165, K562_RS05765 |

### Selected differentially regulated genes at acidic pH in the SCFM-FeZn

Since the genome sequence of *B. cenocepacia* K56-2 was not available in KEGG and the KEGGREST enrichment did not consider genes with different proteins IDs from *B. cenocepacia* J2315, the DE genes list was also analyzed manually. We identified groups of DE genes within genomic clusters. Some genes upregulated at acidic pH were annotated with functions in phosphate metabolism, pili formation, and bacterial chemotaxis (Supplemental Table S1). The 16 upregulated genes associated with phosphate metabolism were part of three gene clusters. In one cluster, a two-component regulator was upstream of three regulatory genes of the phosphate regulon (*phoR*, *phoB* and *phoU)* with all being upregulated, as well as three of the four associated with phosphate specific transport (*pst*) genes (*pstB, pstA* and *pstS*) (43,44). Nine other upregulated genes were part of two clusters encoding for proteins involved in phosphonate (*phn*) transport and metabolism (45,46). Six genes involved in cable pili regulation and structure (*cblRSDCAB*) were also upregulated. Two of these are regulatory genes (*cblR*, *cblS*) and the other four encode proteins involved in pili structure (*cblA, cblB*) and assembly (*cblD*, *cblC*) (47). The structural genes, *cblA* and *cblB*, were among the most highly upregulated genes (27 and 31-fold) at acidic pH. In addition, five genes predicted to be involved in bacterial chemotaxis were also upgregulated (47,48).

Genes involved in oxidative stress response were also upregulated at acidic pH. This includes the transcriptional activators: *soxR*, and the hydrogen peroxide-inducible activator (presumably *oxyR*); a glutathione *S* transferase, two peroxidases, and a catalase. There was increased expression of two genes in oxidative phosphorylation: one encoded part of complex I (K562_RS07885) and another involved in complex III (K562_RS23625).

We found decreased expression of genes associated with ornibactin biosynthesis and uptake, phenylacetate degradation, type III secretion, and cepacian exopolysaccharide production. Ornibactin is a siderophore produced by *B. cenocepacia* under iron-limiting conditions (49,50). All the known genes required for the biosynthesis, export and import of ornibactin are in the same putative operon. Nine of these genes were downregulated at acidic pH. Twelve genes involved in phenylalanine degradation by the phenylacetic acid (*paa*) pathway were downregulated at acidic pH (51,52). These genes are organized in three different putative operons across the *B. cenocepacia* genome(53). Cepacian is an extracellular polysaccharide produced by *Burkholderia* sp. (54). The 21 genes predicted to be involved in cepacian synthesis are organized in two gene clusters: *BceI* and *BceII* (55). Of these genes, 18 had decreased expression at acidic pH. Also, 15 out of the 17 genes associated with type III secretion assembly were downregulated. Genes encoding for proteins involved in the assembly of the type III secretion systems are conserved in multiple *Burkholderia* sp. (56,57). Lastly, we found six genes annotated as cytochrome *c* downregulated at acidic pH. Cytochrome *c* is a periplasmic protein that provides electrons to the complex IV of oxidative phosphorylation.

### B. cenocepacia culture supernatant metabolomics

To further characterize metabolic changes in response to acidic pH in a model of the CF nutritional environment, supernatants of *B. cenocepacia* cultures were analyzed by LC-MS. Metabolites were identified by mass-to-charge ratio (*m*/*z)* and retention time match to a library of authentic standards of an in-house library of common metabolites. Compounds of interest were validated by MS/MS against library standards.

In total, 18 compounds with abundance difference between pH values were detected (fold change > 1.5 and *p*-values < 0.05) (Table 2). Nine compounds were increased in supernatants of cells grown at acidic pH. These included intermediates from central metabolism (pyruvate and fumarate) and from nucleic acid metabolism (adenine, cytosine, L-dihydroorotic acid and hypoxanthine). Other compounds with higher abundance were glutamine, niacinamide and pantoate. Niacinamide is a form of vitamin B_3_ (58), and pantoate is a precursor of pantothenate used in the biosynthesis of coenzyme A (CoA) (59). Glutathione (GSH) was uniquely detected at acidic pH in the SCFM-FeZn supernatants.

**Table 2.** Selected detected compounds in the SCFM-FeZn at acidic pH. *B. cenocepacia* supernatants were analyzed by LC-MS using a hydrophilic interactions chromatography column (HILIC) in positive and negative mode. Compounds with a corrected *p-*value < 0.05 and fold change > 1.5 are shown.

| **Differentially detected compounds** |  |  |  |  |
| --- | --- | --- | --- | --- |
| **Compound** | **Fold Change** |  | **Degree of confidence** | **Corrected *p*-value** |
|  | **Positive mode** | **Negative mode** |  |  |
| 2'-Deoxyadenosine | -2.35 | - | * | 1.52E-06 |
| Adenine<sup>+</sup> | 2.27 | - | * | 9.66E-05 |
| Cytosine | 2.34 | - | * | 1.03E-07 |
| Hypoxanthine<sup>+</sup> | 2.33 | - | * | 4.87E-08 |
| L-Glutamine | 3.76 | 3.62 | ** | 2.57E-10 |
| L-Phenylalanine<sup>+</sup> | -2.84 | -1.51 | ** | 2.15E-16 |
| L-Tryptophan<sup>+</sup> | -2.12 | -1.54 | ** | 3.86E-13 |
| cis-Aconitic acid | - | -2.32 | ** | 6.21E-11 |
| Citric acid/Isocitrate | - | -2.96 | ** | 9.91E-13 |
| Fumaric acid | - | 1.5 | ** | 2.07E-07 |
| Cysteine-S-sulfate | - | -9.6 | ** | 3.15E-13 |
| L-Dihydroorotic acid | - | 10.64 | * | 2.85E-14 |
| Niacinamide<sup>+</sup> | - | 3.61 | * | 8.64E-14 |
| Pantoic acid | - | 1.6 | * | 2.00E-05 |
| Phenylacetic acid | - | -2.88 | * | 1.67E-09 |
| Pyruvic acid<sup>+</sup> | - | 14.05 | * | 2.74E-14 |
| **Uniquely detected in media at either pH** |  |  |  |  |
| **Compound** | **SCFM-FeZn pH** |  | **Degree of Confidence** | **Corrected *p*-value** |
| Glutathione (GSH) | 5.50 |  | * | 9.19E-06 |
| Phosphoenol pyruvic acid | 6.80 |  | * | 5.95E-09 |

(+) Compounds present in media blank;
(\*) Compounds with mean retention time (RT), *m/z* match to reference standard
(\*\*) Compounds with mean retention time (RT), *m/z* and MS/MS fragmentation match to reference standard

Seven compounds were decreased in supernatants of cells grown at acidic pH. Cysteine-*S*- sulfate had the greatest fold decrease (9.5-fold). Interestingly, the aromatic amino acids phenylalanine and tryptophan were decreased, as well as phenylacetate, an intermediate of phenylalanine degradation (60). Deoxyadenosine, and two TCA cycle intermediates (citrate and cis-aconitate) were also decreased. Phosphoenolpyruvate was detected only at pH 6.8.

We created a list of 49 compounds of interest for a targeted analysis of cultured supernatants. This list included secondary metabolites produced by *B. cenocepacia*, and intermediates and precursors of aromatic amino acid metabolism. Compounds with *m/z* matches to targeted list are shown in Supplemental Table S2. A compound consistent to *m/z* values consistent with salicylic acid was increased at acidic pH, while 4-hydroxy-2- heptylquinoline (HHQ), fragin and 2,5-diisopropylpyrazine were decreased. HHQ is a quorum sensing signal molecule, fragin is an antifungal compound, and 2,5-diisopropylpyrazine is a volatile cyclic compound (61–63). A compound consistent with the *m/z* of ornibactin was uniquely detected in supernatants of cells grown at neutral pH. Further characterization of these compounds using authentic standards and MS/MS fragmentation is required to validate their identity.

### Aromatic amino acids metabolism is dysregulated at acidic pH

Comparison of metabolomic and transcriptomic analyses, lead us to identify correlations between DE genes and changes in the levels of compounds from related metabolic processes. Twelve genes involved in phenylalanine degradation by the phenylacetic acid (paa) pathway were downregulated at acidic pH, and phenylalanine, phenylacetic acid and tryptophan had decreased relative abundance in supernatants of *B. cenocepacia* grown at acidic pH. We hypothesized that *B. cenocepacia* uses alternative pathways for phenylalanine consumption at acidic pH. The phenylalanine, tyrosine and tryptophan KEGG metabolic maps were used to manually map DE genes involved in the metabolism of these three amino acids at acidic pH. Two genes involved in tyrosine degradation had decreased expression at acidic pH (K562_RS15355, K562_RS15360). These genes encoded for proteins responsible for the degradation of homogentisate to fumarate. One gene involved in tryptophan metabolism had increased expression at acidic pH (K562_RS02070/ *trpE*). This gene encodes for a subunit of athranilate synthase, which converts chorismate to anthranilate. DE genes and detected metabolites related to aromatic amino acid metabolism are shown in Figure 4.

**Figure 4.**
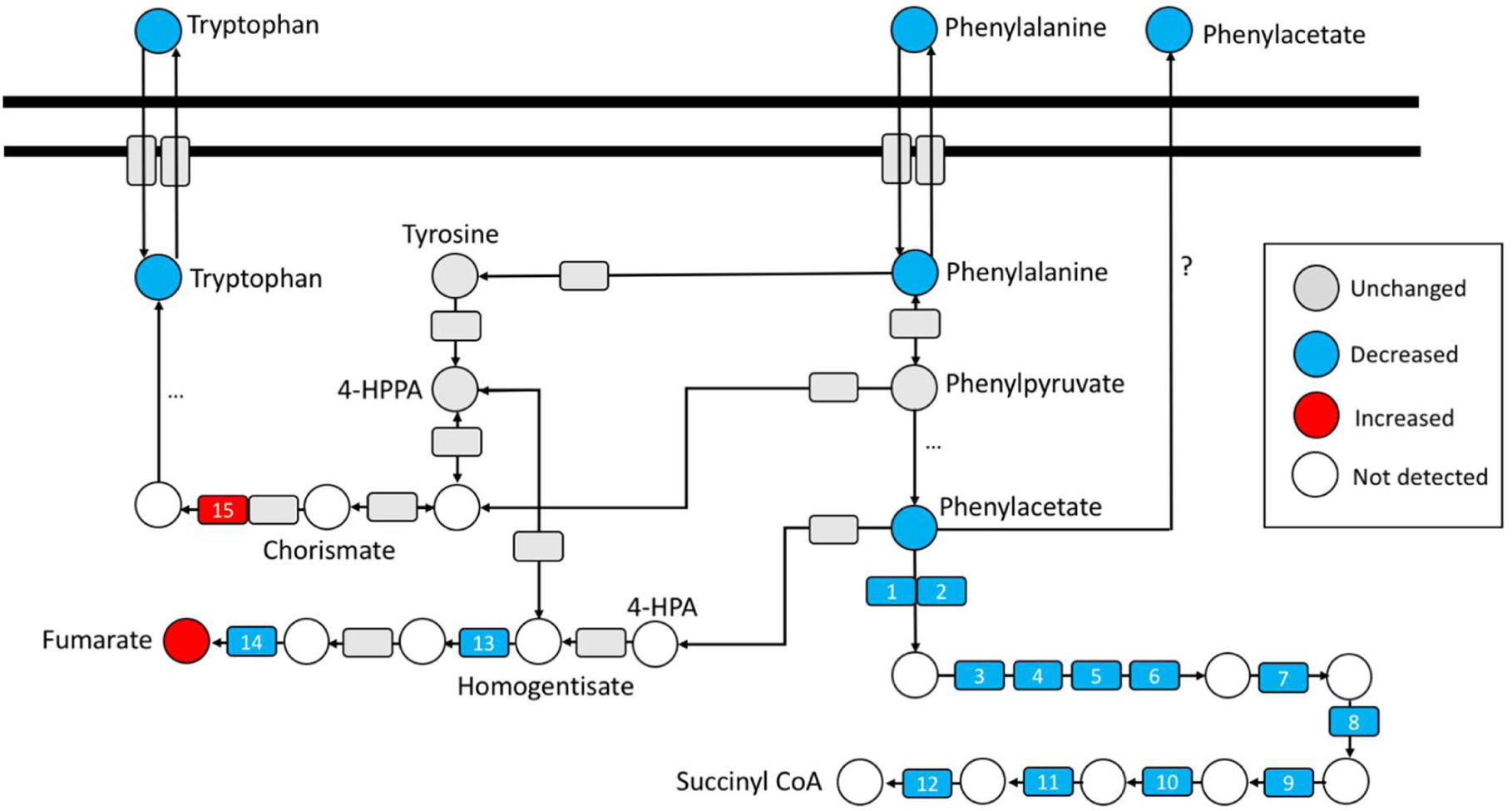
*B. cenocepacia* K56-2 aromatic amino acid metabolism is impacted by acidic pH in SCFM-FeZn. Representation of changes in aromatic amino acids metabolism. Genes are represented as boxes, and metabolites as circles. Gene numbered from 1-12 are involved in the paa pathway, 14-15 are involved in homogentisate degradation, and 15 synthesis of athranilate. Red color indicates increase or upregulation, blue color indicates decrease or downregulation, gray color indicates no difference in expression or production, and white circles represents metabolites that were not detected in the supernatants. Representation includes metabolites with an absolute fold change > 1.5 and corrected *p*-value < 0.05, and differentially expressed genes with an absolute fold change >2 and corrected *p*-value < 0.05. Transcriptomics data represents DE genes from four biological replicates (n=4) of *B. cenocepacia* cells grown at acidic and neutral pH in the SCFM- FeZn. Metabolites data represents mean fold change values from eight biological replicates (n=8) of metabolites detected in *B. cenocepacia* SCFM-FeZn supernatants at acidic and neutral pH.

### B. cenocepacia ΔtrpE growth at acidic pH in the SCFM-FeZn

Increased expression of *trpE* is described in other pathogenic bacteria grown at acidic pH or recovered from macrophages (13,64,65). To assess whether *trpE* contributes to *B. cenocepacia* enhanced growth at acidic pH, we generated a *trpE* deletion mutant in *B. cenocepacia* K56-2 *(ΔtrpE*). *B. cenocepacia* wildtype (WT) and *ΔtrpE* strains were cultured in MH, and SCFM-FeZn at pH 5.50 and 6.80 (Figure 5A). In MH broth, *ΔtrpE* cells reached ∼25% lower cell density compared to the wildtype. In the SCFM-FeZn at both pH values the *ΔtrpE* strain exhibited biphasic growth. At acidic pH, *ΔtrpE* cells entered the second phase of this growth later compared to neutral pH.

**Figure 5.**
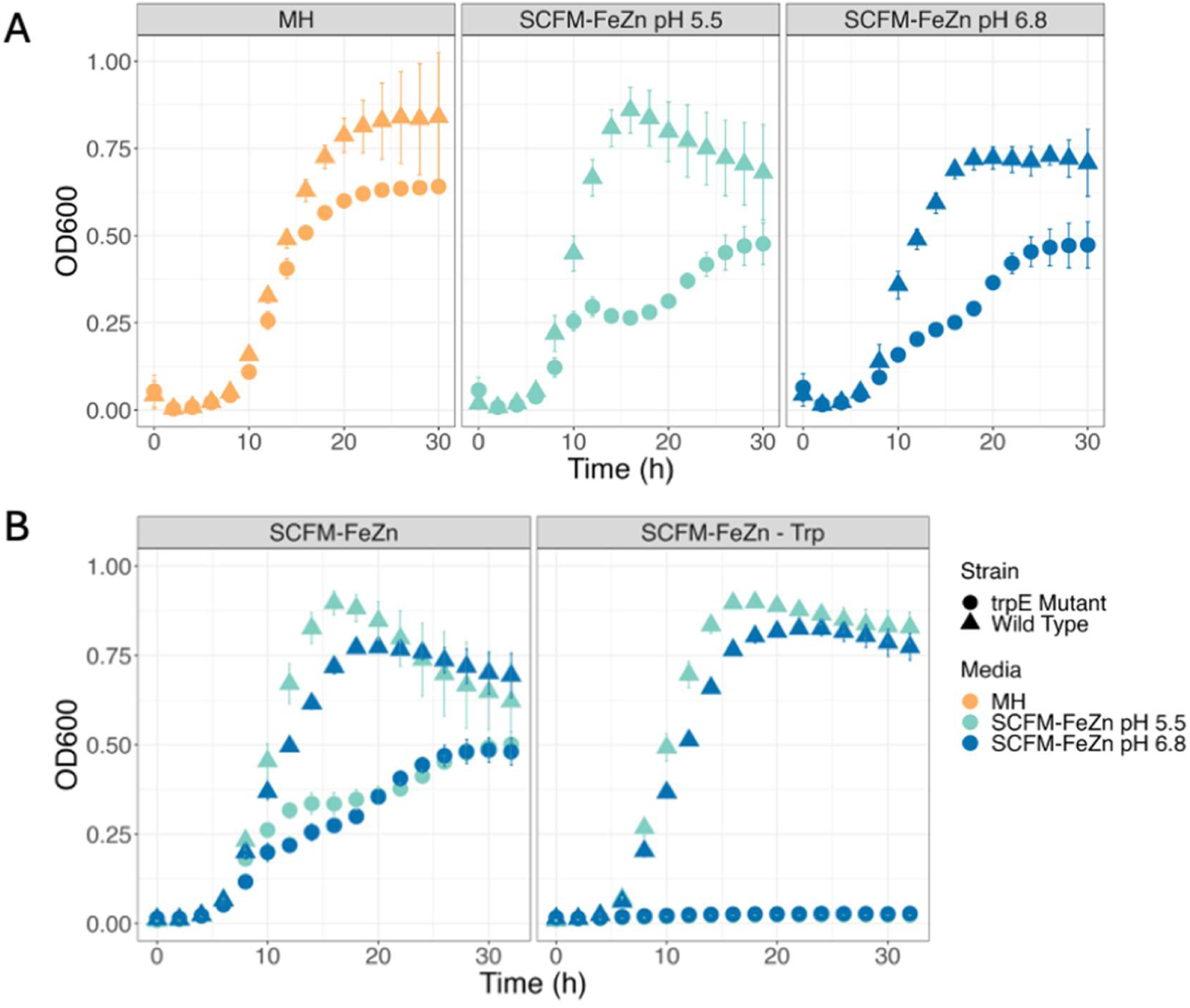
*B. cenocepacia ΔtrpE* mutant growth in MH, SCFM-FeZn at acidic (5.50) and neutral (6.80) pH. **(A)** *B. cenocepacia* WT and ΔtrpE were grown in 100 µl of MH and SCFM- FeZn at pH 5.50 and pH 6.80 (Initial OD_600_ = 0.02). Growth at 37°C was monitored for 32 hours measuring OD600 every two hours. **(B)** *B. cenocepacia* WT and *ΔtrpE* were grown in 100 µl of SCFM-FeZn and SCFM-FeZn without tryptophan (-Trp), at acidic (5.50) and neutral (6.80) pH (Initial OD_600_ = 0.02). Growth at 37°C was monitored for 32 hours measuring optical density (OD600) every two hours. Measurements were taken from six replicates (n=6). Error bars represent the standard deviation between individual measurements (SD).

Deletion of *trpE* in other bacteria resulted in tryptophan auxotroph strains (66). The tryptophan content of the SCFM-FeZn is 10 µM. *B. cenocepacia* WT and *ΔtrpE* strains were grown in the SCFM-FeZn with and without tryptophan (Figure 5B). The *ΔtrpE* strain was unable to grow in the SCFM-FeZn media lacking tryptophan at either pH, while there was no difference in the WT strain growth in either medium. This result confirms that *B. cenocepacia ΔtrpE* is a tryptophan auxotroph.

### B. cenocepacia ornibactin production is decreased at acidic pH

Lastly, *B. cenocepacia* transcriptomics analysis showed decreased expression of genes involved in ornibactin biosynthesis and transport, and correspondingly a compound consistent with *m/z* of ornibactin was detected only in supernatants of cells grown at pH 6.8. The ornibactin gene cluster is composed by fifteen genes of which 8 encode biosynthetic enzymes, 6 encode transport and utilization proteins, and 1 encodes the transcriptional regulator, OrbS (50). Supplemental Figure 4A shows a diagram of the ornibactin biosynthesis pathway and transport, and the genes that were transcriptionally downregulated at acidic pH in SCFM-FeZn. Ferric iron (Fe^3+^) is the predominant oxidation state of iron at neutral pH, and bacteria typically use siderophores such as ornibactin for uptake. In contrast, ferrous iron (Fe^2+^) is the predominant oxidation state at acidic pH, and is acquired using specific ferrous iron transporters (67,68). Thus, siderophore uptake systems are expected to be downregulated at acidic pH, and siderophores to be more abundant at neutral pH.

## Discussion

*B. cenocepacia* encounters acidic conditions in the CF lung sputum and in the phagolysosome of macrophages. Our aim was to characterize the bacterial physiological responses at acidic pH in the CF nutritional context. Interestingly, *B. cenocepacia* maintained a near neutral internal pH (intracellular pH) when exposed to acidic extracellular pH but not to alkaline extracellular pH (Figure 2). The intracellular pH was 7.51 at neutral extracellular pH and is similar to that reported for Gram-negative bacteria with neutral intracellular pH (27,38,69). Similar to *B. cenocepacia*, the intracellular pH of *E. coli* and *P. putida* upon exposure to pH 5.5 remained near neutral (7.0 - 7.5) (27,69). However, at alkaline extracellular pH (8.5), the intracellular pH of *E. coli* is close to 8.50, while that of *P. putida* intracellular pH remains close to neutrality (27). The difference in intracellular pH homeostasis was attributed to lower outer membrane permeability in *P. putida* compared to *E. coli*. Our data suggests that *B. cenocepacia* internal pH homeostasis mechanisms are better adapted to acidic environments compared to alkaline environments. A common mechanism used by bacteria to maintain a neutral internal pH is the expression of carbonic anhydrase, which catalyzes the interconversion of CO_2_ and water to bicarbonate ions and protons (70,71). The carbonic anhydrase (K562_RS19195) expressed in *B. cenocepacia* at acidic pH may function to consume protons (Supplemental Table S1).

*B. cenocepacia* grew faster at acidic pH in SCFM-FeZn and this growth neutralized the media. The response of the bacterium to acidic pH is similar to that described in other Gram-negative bacteria (27,38,69), (42), although some molecular mechanisms and pathways dysregulated in *B. cenocepacia* in the CF nutritional environment have not been described previously. For example, Gram-negative bacteria release bases into the media to increase extracellular pH., In the case of *H. pylori*, ammonia is released from urea while *E. coli* and *Salmonella tyintracellular pHmurium* utilize decarboxylases to release bases such as cadaverine from amino acids (72–74). Interestingly, none of the three decarboxylases (K562_RS08585, K562_RS26105, K562_RS05190) that were expressed in *B. cenocepacia* at acidic pH appear to be homologs of the most commonly expressed enzymes reported in the literature: lysine, arginine, and glutamine decarboxylases (74,75). Other enzymes whose transcripts were more abundant at acidic pH include an amidase (K562_RS14320) and two amidohydrolases (K562_RS30240, K562_RS30265). Although the substrates of these enzymes are unknown, they are potentially used by *B. cenocepacia* to modulate the pH of the CF nutritional environment.

The metabolomics study revealed compounds that are associated with mechanisms to increase extracellular pH (Table 2). For example, hypoxanthine, the product of adenine deamination, was increased in supernatants of cells grown at acidic pH. Ammonia released from this deamination would raise internal pH (76). Glutamine is not a component of SCFM- FeZn but was increased in supernatants at acidic pH. Some Gram-negative enteric bacteria utilize a glutaminase-dependent acid resistance mechanism (77,78). In *E. coli*, this system imports glutamine for deamination in the cytoplasm to release ammonia. In contrast, *Mycobacterium tuberculosis* secretes glutamine into media supernatants and host cells phagosomes (79). The authors of this study suggested this mechanism may mitigate phagosomal acidification. The current data suggest that *B. cenocepacia* may use a similar mechanism.

Another process that we identified in *B. cenocepacia* that is dysregulated at acidic pH in other Gram-negative bacteria is phosphate metabolism. In *B. cenocepacia*, the expression of 16 phosphate metabolic genes was increased at acidic pH (Supplemental Table S1). In other bacteria, these genes are part of the Pho regulon (46,80) and expression is associated to acidic pH adaptation, pathogenicity, and antimicrobial resistance (15,43,81,82) (83). In *P. aeruginosa,* induction of the Pho regulon is associated with biofilm formation, cytotoxicity and motility. Expression of PhoBR-regulated *pst* genes enhances attachment of *E. coli* to host epithelial cells (84) and, in *S. typhimurium*, the Pho regulon protects against acid stress and is associated with bacterial membrane modifications that increase polymyxin resistance (15,85). A role for the Pho regulon in adaptation to acidic stress has not been demonstrated in *B. cenocepacia*.

The increased expression of genes associated with the oxidative stress response in *B. cenocepacia* at acidic pH (Supplemental Table S1) is consistent with what has been reported in other bacteria (17,42). More specifically, acidic stress results in the formation of reactive oxygen species (ROS), inducing genes involved in the oxidative stress response as a detoxification mechanism. Drevinek *et al.* (2008) found that *B. cenocepacia* had increased expression of genes associated with oxidative stress response and iron uptake when grown in media supplemented with sputum from CF patients (86). Similarly, GSH and pyruvate found in cultured supernatants of *B. cenocepacia* grown at acidic pH are associated with stress response (87,88). Smirnova et al. (2012) showed that *E. coli* cells exposed to acidic and alkaline pH increased the secretion of glutathione into the media. Interestingly, these authors observed increased GSH secretion in a strain constitutively expressing Pho regulators. In *E. coli* increased abundance of extracellular pyruvate enhanced acid resistance through activation of RpoS, a regulator of bacterial stress response (89). Also, accumulation of pyruvate inside *E. coli* cells conferred acid resistance.

Importantly, several groups of genes dysregulated in the CF nutritional environment at acidic pH in *B. cenocepacia* encode systems that have not yet been associated with an acid response in other bacteria. Six genes required for cable pili formation had increased expression at acidic pH (Supplemental Table S1). Previous research on *B. cenocepacia* transcriptional response of *cbl* genes in M9 minimal media at acidic and alkaline pH found the highest expression of these genes at pH 6.0, and the lowest expression at pH 5.0 (90). The differing media conditions in our study may alter the pH dependence. *E. coli* cable pilus gene expression was also the highest at pH 7.0 and decreased with pH to 5.0 in LB medium (91). In *B. cenocepacia*, cable pili are required for binding and transmigration across epithelial cells (92). The increased expression of cable pili proteins is consistent with enhanced attachment and invasion of airways epithelia in acidic sputum.

Cepacian is the most common exopolysaccharide produced by bacteria of the *B. cepacia* complex (Bcc) (55,93). The 21 genes associated with cepacian biosynthesis are organized in two gene clusters: *BceI* and *BceII* (54,55). In our dataset, 18 of these genes had decreased expression, suggesting that *B. cenocepacia* produces less cepacian at acidic pH in the SCFM- FeZn (Table S2). Cepacian production is one of the main differences between mucoid and non-mucoid Bcc isolates. Mucoid isolates have been associated with increased persistence and mortality in mice models and decreased recognition by immune cells (94,95). However, Zlosnik *et al.* (2011) showed that patients infected with non-mucoid isolates had more rapid decline of lung function compared to patients infected by mucoid isolates (96). Also, ceftazidime and ciprofloxacin caused conversion of mucoid isolates into non-mucoid. The acidic pH of sputum may be associated with decreased exopolysaccharide production and conversion to the non-mucoid extracellular pH. Cepacian protects against toxic levels of metal ions and cationic antimicrobial peptides as well as other environmental factors (97,98). Thus, decreased expression of cepacian at acidic pH may make *B. cenocepacia* more susceptible to some antibiotics. Nonetheless, *B. cenocepacia* is generally less susceptible to antibiotics at acidic pH, suggesting alternative resistance mechanisms at acidic pH (26).

Integration of transcriptomics and metabolomics data suggests that *B. cenocepacia* phenylalanine and tryptophan consumption increases at acidic pH. We observed decreased expression of twelve genes involved in the phenylacetate (paa) degradation pathway (Table S2), and phenylalanine and phenylacetate were decreased in supernatants of cells grown at acidic pH (Table 2). The paa degradation pathway is a common pathway for aerobic degradation of phenylalanine into the TCA cycle intermediates (99,100). In *B. cenocepacia*, *paa* genes are distributed in three putative operons across the genome: one gene cluster includes *paaABCDE*, another one *paaFZJGIK1*, and the third cluster *paaHK2* (51,53). Hunt *et al*. (2004) found that a Δ*paaE B. cenocepacia* mutant was unable to survive in a rat model (101). In *C. elegans*, infection with *paaA* and *paaE* mutants resulted in increased host survival compared to the wild type, while *paaZ* and *paaF* caused decreased host survival (102). Mutants resulting in accumulation of paa-CoA produced in the first step of the pathway are hypothesized to decrease pathogenesis, while accumulation of other metabolites in the pathway enhance it. Interestingly, comparison of the transcriptomes of *B. cenocepacia* clinical isolates found decreased expression of *paa* genes in isolates recovered from long term infections (103). CF patients with infective exacerbations have lower airway pH than those clinically stable (9). Perhaps during prolonged infection, the pH of sputum decreases and *B. cenocepacia* uses alternative pathways for phenylalanine metabolism.

Only one gene in aromatic amino acid metabolism, *trpE,* was observed to be upregulated at acidic pH. TrpE is a subunit of anthranilate synthase that transforms chorismate into anthranilate (Figure 4). Chorismate is a central metabolite for phenylalanine, tyrosine and tryptophan biosynthesis and anthranilate biosynthesis is the first committed step in tryptophan biosynthesis. Increased expression of *trpE* was reported in *H. pylori* exposed to pH 5.0, and *Listeria monocytogenes* cells recovered from macrophages (13,65). In *M. tuberculosis*, *trpE* deletions resulted in decreased survival in macrophages and tryptophan auxotrophy (64,66,104). These studies suggest that *trpE* expression plays a role in bacterial survival to acidic pH and colonization of the host. In our assays, *ΔtrpE* was a tryptophan auxotroph and had decreased growth in the SCFM-FeZn at both pH values compared to wildtype (Figure 5). We hypothesize that the *ΔtrpE* strain would have decreased survival in macrophages as well. Most studies have attributed the decrease in pathogenesis of *trpE* mutants to tryptophan limitations inside the host (66,104). In *P. aeruginosa* and *Ralstonia solanacearum*, anthranilate is a precursor of quorum sensing molecules (105,106). Interestingly, the anthranilate produced by *R. solanacearum* inhibits hyphae formation and sexual mating of the fungus *Sporisorium scitamineum*. Increased production of anthranilate from the induction of *trpE* expression in *B. cenocepacia* may serve as a precursor for secondary metabolites in addition to tryptophan.

The compounds identified via targeted metabolomics of culture supernatants included metabolites detected in a previous metabolomics analysis of *B. cenocepacia* grown in SCFM2 (107). Similar to the latter study, we found two compounds with *m/z* corresponding to fragin and 2,5-diisopropyl-pyrazine in the SCFM-FeZn. In contrast, we detected a compound with an *m/z* value corresponding to ornibactin only at neutral pH in SCFM-FeZn. The authors of this study only detected ornibactin in LB medium and not in SCFM2. Unlike SCFM-FeZn, SCFM2 contains bovine mucin, salmon sperm DNA, N-acetyl glucosamine and dioleoylphosphatidylcholine. Neve *et al.* (2021), found decreased siderophore production by *P. aeruginosa* in SCFM compared to SCFM2. They attributed this change in iron uptake to increased iron concentrations present in the mucin added to SCFM2 (108), and this rationale may also apply in our study.

Altogether, this study characterized *B. cenocepacia* physiological response to acidic pH in SCFM-FeZn. We elucidated potential mechanisms that contribute to adaptation to the acidic pH of CF sputum. Some of these mechanisms are associated with bacterial persistence, pathogenesis and antimicrobial resistance.

## Materials and Methods

### Measuring the pH of SCFM-FeZn (pH_m_)

To track the changes in media pH during *B. cenocepacia* growth, the fluorescent pH indicator BCECF (2′,7′-bis-(carboxyethyl)-5-(and-6)-carboxyfluorescein) was used. BCECF was calibrated in the SCFM by adjusting pH values from 5.0 to 8.5. SCFM-FeZn (100 μl) at each pH value was supplemented with 0.1 μM BCECF, aliquoted in a 96-well plate and fluorescence measured (λ_ex_ = 500 nm, λ_em_ = 530 nm). A calibration curve of fluorescence plotted against media pH (Figure S1) was used to transform fluorescence measurements into pH values.

SCFM-FeZn with low buffering capacity (10 mM MOPS, 10 mM MES) and high buffering capacity (25 mM MOPS, 25 mM MES) was used in growth assays. The pH of the SCFM-FeZn was adjusted to 5.5 or 6.8, and medium was supplemented with 0.1 μM BCECF, and aliquoted into a 96-well plate. *B. cenocepacia* precultures were set from overnight cultures and each well was inoculated to an initial OD_600_ of 0.02. Cells were incubated for 24 h at 37 °C inside the plate reader (Tecan Infinite PRO 200), fluorescence (λ_ex_ = 500 nm, λ_em_ = 530 nm) and OD_600_ measurements were taken every four hours. Cells were shaken before every measurement.

### Measuring B. cenocepacia K56-2 internal pH (intracellular pH)

The PHP (**pH** indicator for ***P****seudomonas*), developed by Arce-Rodriguez *et al.* 2019, was used to track the internal pH (intracellular pH) of *B. cenocepacia* K56-2 (27). The vector containing the PHP sequence was purchased from Addgene (pS2513-PHP; catalog number: #122590). The PHP gene was cloned into a vector optimized for gene expression in *B. cepacia* (pMLBAD-eGFP; Addgene catalog number: #112918) (28) between the NcoI and XbaI restriction sites using a T4 DNA ligase. *E. coli* DH5α cells were transformed with ligation product, plated in LB agar supplemented with 50 μg/ml trimethoprim and incubated at 37 °C. Trimethoprim resistant colonies were screened using colony PCR. The pMLBAD-PHP vector was then purified and presence of insert was confirmed by sequencing.

The pMLBAD-PHP vector was transformed into *B. cenocepacia* by electroporation and plated on SOC plates supplemented with 100 μg/ml trimethoprim. To assess the pH-sensitive spectrum of PHP, *B. cenocepacia* cells carrying the pMLBAD-PHP plasmid were grown in overnight cultures and precultures of Mueller Hinton (MH) supplemented with 2% L-arabinose and 100 μg/ml trimethoprim. Cells were harvested from precultures at an OD_600_ between 0.4 and 0.6 by centrifugation (4000 × *g* for 15 min). Cells were then suspended in 200 μl M9 minimal media at pH values ranging from 5.0 to 8.5 supplemented with membrane uncouplers (50 mM sodium benzoate and 50 mM methylamine HCl) to a final OD_600_ of 10. M9 media was supplemented with 50 mM of the following buffers depending on the targeted pH of the media: MES (pH 5.0, 5.5 and 6.0), PIPES (pH 6.5 and 7.0), MOPS (pH 7.5), and TAPS (pH 8.0 and 8.5). Cell suspensions were added to a 96-well plate and the excitation spectrum was measured from 350 to 500 nm (λ_em_ = 515 nm). All fluorescence measurements were taken with a Tecan Infinite PRO 200.

To build a PHP calibration curve, *B. cenocepacia* harboring pMLBAD-PHP was grown as previously described. Cells were harvested and suspended in 200 μl of M9 media at different pH values supplemented with membrane uncouplers to a final OD_600_ of 10. Cell suspensions were added to a 96-well plate and fluorescence was measured (λ_ex_ = 400 and 478 nm, λ_em_ = 515 nm). The calibration curve was built by calculating the fluorescent ratios between the two excitation wavelengths (400 nm/478 nm) and plotting them against media pH.

To measure the intracellular pH of *B. cenocepacia* after shifts to acidic and alkaline pH, cultures of *B. cenocepacia* harboring pMLBAD-PHP were aliquoted into Falcon tubes, suspended in 200 μl of M9 medium at pH 7.0 and incubated at room temperature for five minutes (final OD_600_ = 10). Cells were pelleted again and resuspended in 200 μl of M9 at pH 5.5, 7.0 or 8.5. Fluorescence was measured every 20 s for ten minutes. To assess the intracellular pH in SCFM-FeZn, cells were harvested and suspended in M9 at pH 7.0 for five minutes, then pelleted again and resuspended in 200 μl of SCFM-FeZn at pH 5.5 and 6.8. Buffered M9 and SCFM-FeZn supplemented with 50 mM of sodium benzoate and 50 mM methylamine HCl were used as controls. Fluorescent ratios (400 nm/478 nm) were used to calculate the intracellular pH of the cells using a PHP calibration curve.

### Transcriptomics analysis

*B. cenocepacia* K56-2 overnight cultures were inoculated from eight individual colonies and incubated for 16 h at 37 °C. For precultures, 5 ml of MH broth were inoculated with 250 μl of overnight cultures, incubated for 4 – 6 h, and 1 ml of each preculture was spun down and suspended in 0.85% saline solution. The saline resuspension was used to inoculate 5.5 ml SCFM-FeZn at pH 5.5 or 6.8 to an initial OD_600_ of 0.02. Inoculated SCFM-FeZn was incubated at 37 °C (50 rpm) to an OD_600_ of 0.4 - 0.6 (13 h). *B. cenocepacia* K56-2 cells were harvested by centrifugation (4000 × *g* for 10 min), suspended in 1 ml RNA*later* stabilization solution (Invitrogen^TM^) and left at 4 °C for 24 h. Cells were then pelleted by centrifugation, RNA*later* was removed, and pellets were flash frozen in liquid nitrogen and stored at -70 °C. Culture supernatants were spun down (16000 × *g* for 10 min), flash frozen in liquid nitrogen and stored at -70 °C.

Total RNA from *B. cenocepacia* K56-2 cells was extracted using RiboPure^TM^-Bacteria Kit (Invitrogen^TM^) according to manufacturer’s instructions. Contaminating genomic DNA was removed from RNA samples using with Turbo^TM^ DNase (Invitrogen^TM^). Two μl of SuperaseIn RNase inhibitor (Invitrogen^TM^) was added to each sample before DNase treatment and 1 μl after treatment. RNA quality was assessed using a 2100 Bioanalyzer (Agilent), and quantity was determined using Quant-iT (ThermoFisher). Ribosomal RNA depletion and cDNA library construction was performed using the QIAseq FastSelect 5S/16S/23S kit (Qiagen) and KAPA RNA HyperPrep Kit (Roche), respectively. Single-end 150 base pair sequencing reads were generated on the Illumina HiSeqX platform. Raw sequence quality control was assessed using FastQC. Reads were aligned to the reference genome using STAR aligner (version 2.7.9a) (29).

*B. cenocepacia* K56-2 reference genome was downloaded from RefSeq (GCF_014357995.1) (30). Summary of the reads aligned per gene was generated with htseq-count (version 0.11.2) (31). DESeq2 R package (version 1.26.0) was used to find differentially expressed genes in cells grown at acidic (5.50) versus neutral pH (6.80) in SCFM-FeZn (32). Genes with more than two-fold change and *p*-value < 0.05 were selected. Since *B. cenocepacia* K56-2 pathway annotations were not available, relevant annotations from a closely related strain, *B. cenocepacia* J2315, were used to perform pathway enrichment analysis.

Pathway annotations for *B. cenocepacia* J2315 were downloaded from www.burkholderia.com (DB version 9.1; 2020-04-20) on Sept 22, 2022. This included all pathway annotations from MetaCyc (33), KEGG (34), and InterPro (35). Pathways assigned to proteins in *B. cenocepacia* J2315 were mapped to proteins in *B. cenocepacia* K56-2 where protein GenBank accession numbers were identical. In addition, KEGGREST (version 1.26.0) was used to access KEGG pathways assigned to *B. cenocepacia* J2315 genes and identical proteins in *B. cenocepacia* K56-2. DE gene lists were used to apply a Wilcoxon rank-sum test to test for enrichment of each pathway.

### Metabolomics analysis

Liquid chromatography mass spectrometry (LC-MS) analysis performed using an Agilent 1290 Infinity II UHPLC in line with an Agilent 6546 Q-TOF with a dual AJS ESI source. Culture supernatants collected as described above in the transcriptomics section were passed through a 0.2 μm PTFE syringe filter. Two µl sample injections were separated using HILIC and c18. For HILIC, an InfinityLab Poroshell 120 HILIC-Z column (100 mm × 2.1 mm × 2.7 μm) was used with a 10 min linear gradient from 90 to 60% Solvent B at 0.25 ml/min. Solvent A was water with 10 mM ammonium acetate, pH 9 and 5 μM medronic acid and Solvent B was 10 mM ammonium acetate, pH 9 in 90% acetonitrile. For c18, a Zorbax Eclipse Plus c18 column (100 mm × 2.1 mm × 1.8 µM) was used with a 16-minute linear gradient from 5 to 100% solvent B at 0.45 ml/min. Solvent A was 0.1% formic acid in water, solvent B was 0.1% formic acid in acetonitrile. MS parameters in negative ionization mode were as follows: capillary voltage, 3500 V; nozzle voltage, 1000 V; drying gas temp, 250 °C; drying gas flow rate, 10 l/min; sheath gas temperature, 300 °C; sheath gas flow rate 12 l/min, nebulizer pressure, 40 psi; fragmentor voltage, 100 V. Parameters for positive ionization mode were the same, except the nozzle voltage, 500 V. MS/MS was collected on selected ions with 10, 20, and 40 V collision energies. Data were collected and analyzed using MassHunter Profinder v. 10.0 and Mass Profiler Professional 15.1 (Agilent Technologies). Compounds were identified by an *m*/*z* and retention time match to authentic standards and compounds of interest were validated by MS/MS against library standards. For compounds not present in the available libraries, identification was based on *m/z* matches alone. Statistical analyses were performed using one-way ANOVA and Tukey’s multiple testing correction. Compounds with a corrected *p*-value < 0.05 and fold change > 1.50 were selected.

### Generation and characterization of B. cenocepacia ΔtrpE

A *B. cenocepacia* K56-2 *trpE* deletion *(ΔtrpE*) was generated using homologous recombination following the protocol of Flannagan *et al.* (36). To generate the pGPI-SceI-*ΔtrpE* mutagenesis plasmid, 600-700 bp upstream (US) and downstream (DS) regions flanking *trpE* were PCR-amplified using the following primers: Fwd_trpE_US (5’ – TTGCACGTACAGGTTGCCGTTGTAGAGGGACAGCGGCGTTTC – 3’), Rev_trpE_US (5’- AACGGCTGACATGGGAATTCCGGGATGGCGACGCTGAC – 3’), Fwd_trpE_DS (5’- GATCCCAAGCTTCTTCTAGATTCGACCGGCAGCGTCTTGT-3’), and Rev_trpE_DS (5’ – GAAACGCCGCTGTCCCTCTACAACGGCAACCTGTACGTGCAA - 3’), and *B. cenocepacia* K56-2 genomic DNA as a template. US and DS regions were amplified using Q5 polymerase and the following thermal cycling parameters: 98 °C for one min followed by 30 cycles of 98 °C for 20 s, 70 °C for 30 s and 72 °C for two min, and a last step of 72 °C for two min. The pGPI-SceI vector was digested with XbaI and EcoRI and US/DS region was inserted using Gibson Assembly reaction (NEB: #E2611S). *E. coli* GT115 cells were transformed by heat shock treatment with 4 µl of the assembly reaction. Transformed cells were recovered for one h in LB shaking at 37 °C before plating on LB agar with 50 µg/ml trimethoprim. Colonies containing the pGPI-SceI-*ΔtrpE* vector were screened using the pGPI-SceI primers Fwd_pGPI_SceI (5’-TAACGGTTGTGGACAACAAGCCAGGG-3’) and Rev_pGPI_SceI (5’- GCCCTACACAAATTGGGAGATATATC - 3’). The resulting plasmid insert was confirmed by sequencing.

The pGPI-SceI-*ΔtrpE* vector was introduced into *B. cenocepacia* K56-2 by conjugation using *E. coli* GT115 carrying the plasmid as the donor strain, and *E. coli* DH5α carrying pRK2013 as a helper strain. *B. cenocepacia* mutants were selected for tetracycline-resistance and trimethoprim susceptibility. Colonies were screened by PCR using the Fwd_*trpE*_DS and Rev_*trpE*_DS primers previously described. To cure the pDAI-SceI-*ΔtrpE* vector from *B. cenocepacia*, colonies with the deletion were grown overnight in LB without antibiotics. The region containing the mutation was amplified and sent for sequencing.

### B. cenocepacia ΔtrpE growth in the SCFM-FeZn

*B. cenocepacia* K56-2 wild type and *ΔtrpE* precultures were grown in MH as previously described. Precultures were spun down and suspended in 0.85% NaCl solution. Cell suspensions were used to inoculate 100 µl of MH and SCFM-FeZn at pH 5.5 and pH 6.8 to an initial OD_600_ of 0.02. The inoculated 96-well plate was incubated at 37 °C for 32 hours inside a plate reader (Tecan Infinite PRO 200). OD_600_ measurements were taken every two hours. Cells were shaken before every measurement. To assess the growth of the *ΔtrpE* strain in the absence of tryptophan, growth of WT and *ΔtrpE* strains was also followed in SCFM-FeZn at pH 5.5 and 6.8 without tryptophan.

## Acknowledgments

This work was supported by a Cystic Fibrosis Canada Research Grant to M.E.P.M. and a four-year fellowship (4YF) to L.D.M. from The University of British Columbia. Reza Falsafi depleted rRNA and constructed the cDNA library. We thank Dr. Miguel Valvano (Queens University, Belfast) for the *B. cenocepacia* strain used in this study and Dr. Silvia Cardona (University of Manitoba) for some of the plasmids used for *B. cenocepacia* mutagenesis.

